# Ratiometric Sensing of Redox Environments Inside Individual Carboxysomes Trapped in Solution

**DOI:** 10.1101/2022.03.17.484789

**Authors:** William B. Carpenter, Abhijit A. Lavania, Julia S. Borden, Luke M. Oltrogge, Davis D. Perez, Peter D. Dahlberg, David F. Savage, W. E. Moerner

## Abstract

Diffusion of biological nanoparticles in solution impedes our ability to continuously monitor individuals and measure their physical and chemical properties. To overcome this, we previously developed the Interferometric Scattering Anti-Brownian ELectrokinetic (ISABEL) trap, which uses scattering to localize a particle and applies electrokinetic forces which counteract Brownian motion, thus enabling extended observation. Here, we present an improved ISABEL trap that incorporates a near-infrared scatter illumination beam and rapidly interleaves 405 and 488 nm fluorescence excitation reporter beams. With the ISABEL trap, we monitor the internal redox environment of individual carboxysomes labeled with the ratiometric redox reporter roGFP2. Carboxysomes widely vary in scattering contrast (reporting on size) and redox-dependent ratiometric fluorescence. Further, we used redox sensing to explore the chemical kinetics within intact carboxysomes, where bulk measurements may contain unwanted contributions from aggregates or interfering fluorescent proteins. Overall, we demonstrate the ISABEL trap’s ability to sensitively monitor nanoscale biological objects, enabling new experiments on these systems.

**TOC Graphic:** 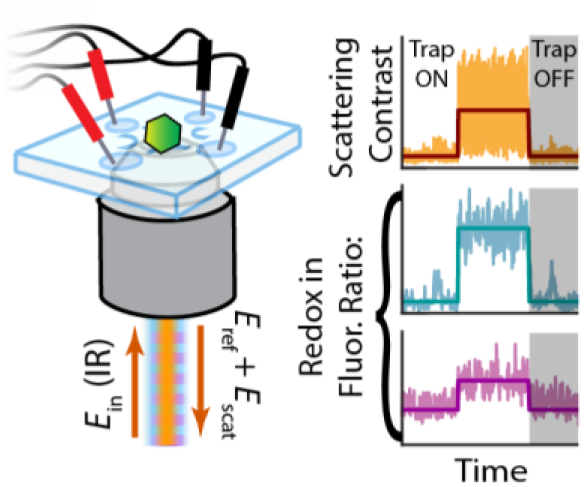

## Main Text

For nanoscale biological objects in solution, Brownian fluctuations dominate their translational dynamics. Due to their stochastic trajectories and fast diffusion, individual objects are commonly immobilized for extended study,^1,2^ which may undesirably perturb them from their native states.^1–5^ Particles on the order of ~100 nm in diameter can be rapidly followed by stage motion,^6, 7^ but this strategy requires a large area passivated surface. To make extended measurements without tethering, Anti-Brownian ELectrokinetic (ABEL) traps have been developed^8^ to apply electrokinetic positional feedback on single fluorescent objects and thus greatly reduce Brownian motion.^9–11^ These traps have been used to directly measure the dynamics of single enzymes,^12^ photosynthetic complexes,^13–16^ and even single organic fluorophores.^17^ Typically, these molecules can be held for seconds, when photobleaching or blinking interrupts continuous positional monitoring.

To overcome the need for fluorescence to estimate position, our lab recently developed the Interferometric Scattering ABEL (ISABEL) trap, which tracks a nanoparticle’s position by its scattering interfered with a local oscillator arising from the back reflection off a water-quartz interface in the sample cell.^18^ The interference between the scattered and reflected light enhances sensitivity by producing a signal that scales linearly with particle polarizability, which may be interpreted as mass for objects with fixed composition.^19–21^ Our initial study demonstrated trapping of gold nanoparticles as small as 20 nm, and that polymer nanoparticles down to 50 nm diameter could be held for more than 30 seconds. Fluorescently labeled particles could also be trapped far beyond the time of photobleaching. Interferometric positional monitoring, independent of fluorescence, opens the door to studying weakly fluorescent biological objects or introducing complex fluorescence excitation protocols.

One such nanoscale biological object is the carboxysome, a proteinaceous microcompartment ~100 nm in diameter, which is responsible for the fixation of CO_2_ into organic carbon in many autotrophic bacteria.^22–25^ The chemoautotroph *Halothiobacillus neapolitanus* contains α-carboxysomes, whose operon encodes 10 proteins that collectively assemble into carboxysomes, including: rubisco large (CbbL) and small subunits (CbbS), a disordered scaffolding protein (CsoS2), carbonic anhydrase (CsoSCA), two pentameric shell protein paralogs (CsoS4AB), three hexamer shell protein paralogs (CsoS1ABC), and a pseudo-hexameric shell protein (CsoS1D).^23,26^ The self-assembled accumulation of rubisco and carbonic anhydrase inside the roughly icosahedral^27^ shell (Fig. 1a) has evolved to create a high local concentration of rubisco and CO_2_ to overcome rubisco’s slow turnover rate and outcompete deleterious oxygenation reactions.^28^ Functional carboxysomes can also be recombinantly grown in *E. coli*,^22^ which aids in inserting fluorescent reporters but also increases the diversity of shapes, sizes, and integrity of the shell (Fig. 1b). This paper reports exclusively on *E. coli*-derived carboxysomes.

**Figure 1.**
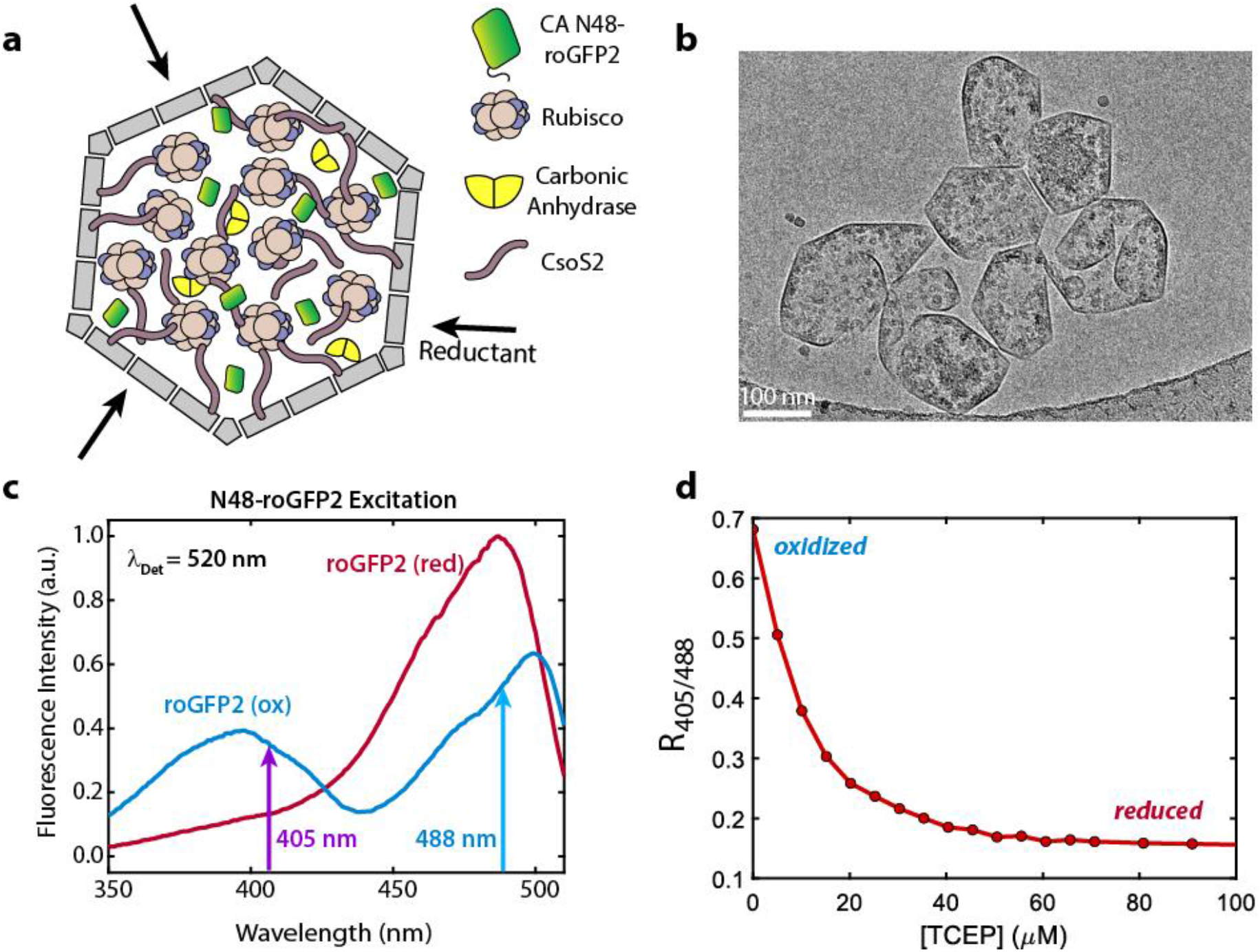
Visualizations of carboxysomes and characteristics of the redox-sensitive GFP mutant roGFP2. (a) The carboxysome consists of a porous proteinaceous shell and the internal cargo rubisco, carbonic anhydrase, and the scaffolding protein CsoS2. roGFP2 is targeted inside the carboxysome using the N-terminal sequence from carbonic anhydrase. (b) A cryo-TEM image of a cluster of α-carboxysomes recombinantly expressed in *E. coli*, demonstrating the variety of shapes, sizes, and integrity. (c) Changes in the fluorescence excitation spectrum of roGFP2 enable ratiometric readout of the redox environment. The fully oxidized spectrum (blue) is bimodal and gives a high fluorescence ratio *R*_405/488_, while the fully reduced spectrum (red) consists of one peak and produces a low fluorescence ratio. The vertical arrows indicate the excitation wavelengths used in this study for ratiometric measurements. (d) Ratiometric fluorescence from roGFP2 decreases when reductant is added.

Structural and simulation studies posit that the proteinaceous shell preferentially allows the bidirectional diffusion of metabolically important species such as HCO3-, and ribulose-1,5-bisphosphate,^29,30^ and therefore is expected to support a distinct chemical environment from the surrounding cytosol.^31, 32^ The protein shell is also thought to establish a distinctly oxidizing redox environment within the carboxysome relative to the known reducing environment of the cytosol.^28, 32–35^ These hypotheses remain unconfirmed, however, due to the lack of direct measurements on selective shell permeability and redox dynamics. Because of the variation of carboxysome size, shape, and integrity, and to mitigate contamination from purification byproducts, it would be highly beneficial to study carboxysomes at the single-particle level. Our goal is to not only trap, but also to sense the redox chemical environment inside individual carboxysomes using a local fluorescent protein reporter, roGFP2, which encodes redox information in its fluorescence excitation spectrum (Fig. 1c). We have genetically targeted approximately 3-15 copies of roGFP2 inside individual carboxysomes (Fig. S1). The ratio of fluorescence brightness from 405 nm and 488 nm excitation is related to the concentration of reducing species in solution (Fig. 1d), and gives a readout that does not rely on GFP copy number.

To enable fluorescence excitation spectroscopy of roGFP2 inside carboxysomes, we have redesigned the ISABEL trap.^18^ In the new configuration (*vide infra*), the scatter illumination beam has been red-shifted to 800 nm in the near-IR to open up the visible region for fluorescence reporters without photobleaching them. As well, we have introduced two rapidly interleaving excitation beams at 405 and 488 nm to measure the fluorescence emission from roGFP2.^36^

In this paper, we directly measure the redox-dependent ratiometric fluorescence of single trapped carboxysomes, where air-oxidized carboxysomes show much more heterogeneous ratiometric fluorescence than reduced carboxysomes. Despite this heterogeneity, we also observe reduction kinetics in carboxysomes after mixing with reductant, in a first step towards measuring shell permeability. Together, these measurements demonstrate the ability of the ISABEL trap to go beyond synthetic nanoparticles to make extended measurements on single biological objects, and that single-particle measurements on individual carboxysomes provide a new avenue for measuring their physical and chemical properties.

Figure 2 shows the quartz microfluidic trapping cell and the optical layout of the ISABEL trap. Nanoparticles in aqueous solution are allowed to diffuse to the center of the cell’s two crossed channels in a region 1.5-2 μm deep (Fig. 2a). When an infrared electric field *E*_i_ is incident on the particle, the back-scattered field *E*_s_ interferes with the quartz-water interfacial reflection, *E*_r_, producing a detected intensity:

**Figure 2.**
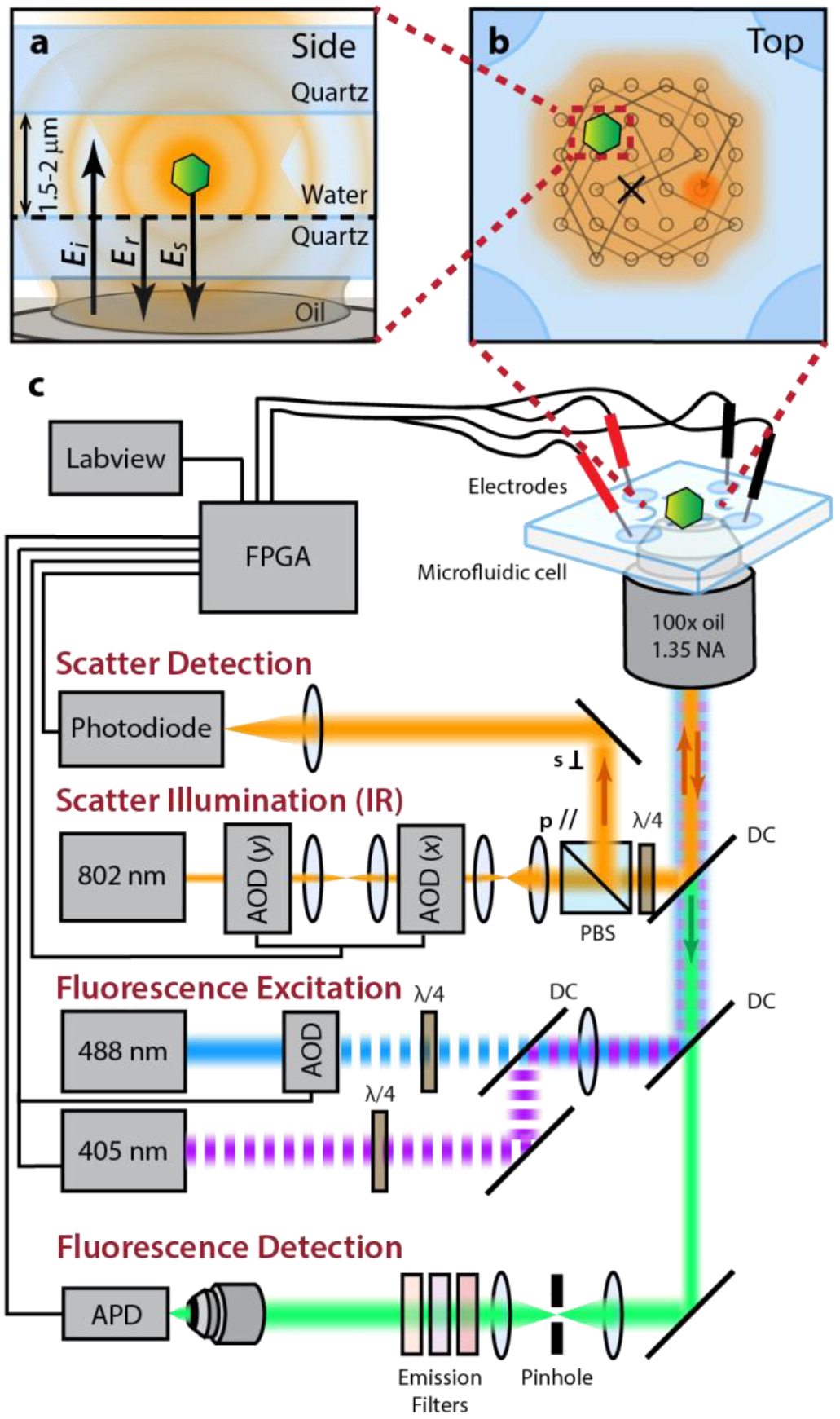
Overview of ISABEL Trap with interleaved fluorescence excitation. (a) A focused incident field *E*_i_ illuminates a carboxysome, which radiates the scattered electric field *E*_s_ that interferes with *E*_r_, the reflection from the quartz-water interface. (b) Top view of the microfluidic trap, where the incident beam is scanned in a 32-spot Knight’s tour pattern. Electrokinetic feedback in two dimensions is applied to the particle to push the object toward the center of the illumination scan pattern (marked “x”). (c) Optical paths of scatter and fluorescence beams, described in more detail in Note S2 and Fig. S2. The scatter illumination beam is deflected by two acousto-optic deflectors (AODs) controlled by the FPGA; it is linearly polarized at the polarizing beam splitter (PBS), then converted to circular polarization with a quarter-wave plate (λ/4) to be back-reflected, converted back to the orthogonal linear polarization, and separated for detection on a photodiode. Position is monitored and feedback voltages are calculated on the FPGA, then applied to the solution with platinum electrodes. Simultaneously, the FPGA digitally modulates two CW fluorescence excitation lasers, alternating each ms. Fluorescence emission spanning 500-570nm is collected on an avalanche photodiode (APD) after spatial filtering with a 75μm pinhole. Detected photons are time tagged on the FPGA and labeled with the identity of the corresponding excitation laser. AOD: acousto-optic deflectors, AOM: acousto-optic modulator, PBS: polarizing beam splitter, DC: dichroic beamsplitter.

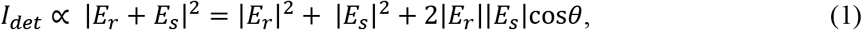

where *θ* is the phase between the reflected and scattered fields. The last term on the right-hand side of Eq. 1 represents the interferometric contribution to the intensity, which is linear in scattered field and thus linear in polarizability and mass for proteinaceous objects.^37^ For small particles, |*E*_s_|^2^ is negligible, and |*E*_r_|^2^ can be obtained by a background measurement when there are no particles in the trapping area. This allows for on-the-fly determination of the absolute fractional scattering contrast

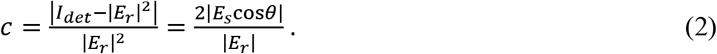

The particle position is detected by the location of the maximum contrast in the sample plane, obtained from a “Knight’s tour” scan pattern of the near-IR beam steered by two acousto-optic deflectors.^9^ Rapid positional feedback forces are applied to the solution via voltages calculated on a Field-Programmable Gate Array (FPGA) and two pairs of platinum electrodes placed at the ends of the crossed microfluidic channels. The particle is directed to the trap center in two dimensions by electroosmosis.

In addition to the IR trapping beam, the second major difference from previous work is the addition of two visible lasers in wide-field illumination for spectroscopic measurements of the trapped object (Fig. 2c). The FPGA digitally modulates each laser power with a 2-ms alternating square wave, so that emitted photons from GFP fluorescence can be separated into two excitation channels (Note S2).

Three simultaneous variables are monitored in time for individual trapped carboxysomes: absolute fractional scattering contrast (Note S3), emission from 405 nm excitation, and emission from 488 nm excitation (Fig 3). A step change in the fractional scattering contrast trace at *t* ≈ 51 s (Fig. 3a, Event i) shows that when trapping turns on, a diffusing particle becomes trapped for >1s, then leaves when the feedback is turned off. For these experiments, we toggled feedback on for 2s and off for 1s to collect statistics from additional single particles, although we can trap carboxysomes for tens of seconds if desired (Fig. S3). The scattering trace of Event i shows a sudden increase in scattering contrast about a mean value (dark red line) determined by a changepoint algorithm described below, with wide fluctuations due to the evolving phase *θ* between *E*_r_ and *E*_s_ as the particle diffuses in the axial direction. Subsequent trapping events display various mean scattering values, indicating a range of particle sizes, including an exceptionally large particle (Event ii), which is likely an aggregate of multiple carboxysomes and is excluded from further analysis. At *t* ≈ 75s (Event iii), a somewhat small object is trapped, which is then replaced by an object of higher contrast (Event iv), since anti-Brownian feedback can be applied to only one object at a time.

**Figure 3.**
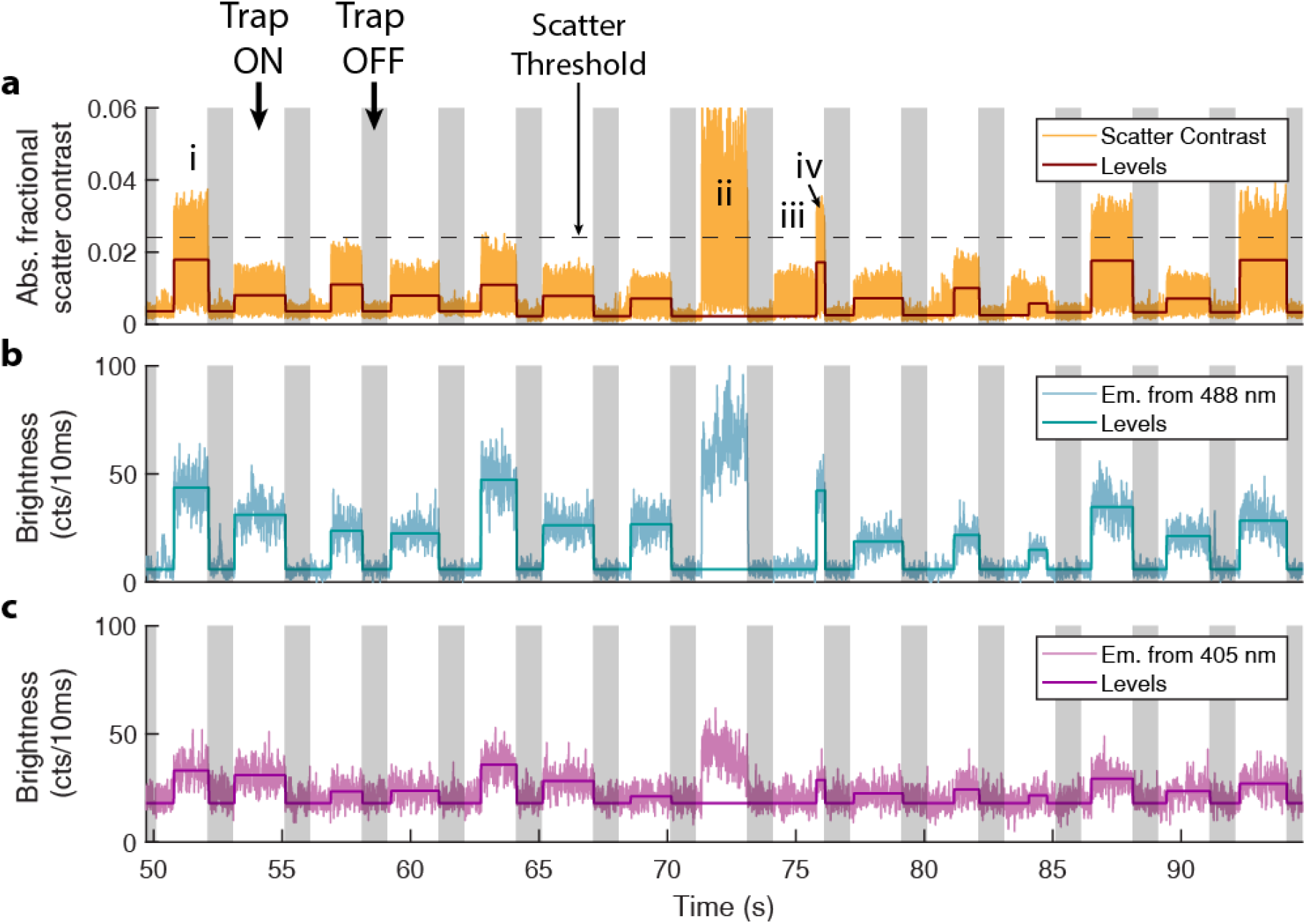
Representative multi-channel time traces from a carboxysome trapping experiment in air-oxidized buffer. When positional feedback is turned on (white regions), single detected particles are held at the trap center until feedback is toggled off (gray regions). Trapped roGFP2-labeled carboxysomes display signal in all three channels, such as in Event i. (a) Absolute fractional scatter contrast trace, with individual measurements for each timepoint (yellow) and average levels for an event (red) determined by the changepoints found on the 488 nm trace (b). Black dashed line indicates the scatter threshold used to reject large aggregates from analysis (Note S3). (b)-(c) 488 and 405 nm excitation channels, respectively, and corresponding average levels determined from a changepoint finding algorithm on the 488 nm trace.

Simultaneously, we monitor fluorescence from each carboxysome via the interleaved 405 and 488 nm excitation (Figs. 3b and 3c, respectively). Low intensity (< 50 W/cm^2^) is necessary at 405 nm to balance the roGFP2 emission rate with the light-induced photoconversionr^38^ of roGFP2 chromophores over extended trapping times (Figs. S3 and S4). In Event i, a steady fluorescence level is present in both channels over the trapping time, but particles are brighter and background is lower in the 488 nm channel. Fluorescence changepoints and mean fluorescence brightness were determined with a changepoint finding algorithm^39^ on the 488 nm excitation trace, which provided changepoints to also find the average levels in the scattering and 405 traces. Like the various scattering levels, the fluorescence traces show a distribution of brightnesses, indicating variation in roGFP2 loading between carboxysomes. The highly scattering object in Event ii is accompanied with high brightness in both channels. The object in Event iii is non-fluorescent, possibly being an emptied carboxysome shell, a protein aggregate that remained after purification, or a dislodged piece of the polyelectrolyte passivation layer (Note S1). Because this particle is non-fluorescent, it is not identified by the algorithm. Conversely, the object in Event iv shows signal in all three channels, implying that it is a carboxysome.

The simultaneously measured levels from scattering and fluorescence provide correlated data from individual trapping events, yielding multidimensional statistics measured from carboxysomes in reducing or air-oxidized buffers (Fig. 4). Figs. 4a-4c show 2D scatter plots from carboxysomes internally reduced by 1 mM TCEP in buffer. In these plots, each point represents the average level found from an individual trapping event, and its color reflects the local density of points.^4^ In the Fig. 4a marginal histogram, the 488 nm fluorescence distribution is centered at ~60 counts/10ms but spans two orders of magnitude. The 2D scatter plot shows correlation between fluorescence levels and fractional scattering contrast, with subpopulations within the spread. The 2D scatter plot in Fig. 4b relating 405 nm brightness to scattering shows similar trends, though with lower brightnesses values centered at ~5 counts/10ms. The scattering constrast histogram (Fig. 4c right) appears bimodal, peaked at 0.008 and 0.015. In contrast, cryo-TEM imaging reveals that the distribution of carboxysome diameters is unimodal (*μ* ± *σ* = 141 ± 31 nm, Fig. S5). Because of the approximate doubling in scattering contrast and unimodal diameter distribution, the higher contrast peak likely arises from trapped carboxysome dimers. The monomer and dimer populations are not readily separable in a 1D measurement, though they are better separated with our multi-dimensional correlated measurements.

**Figure 4.**
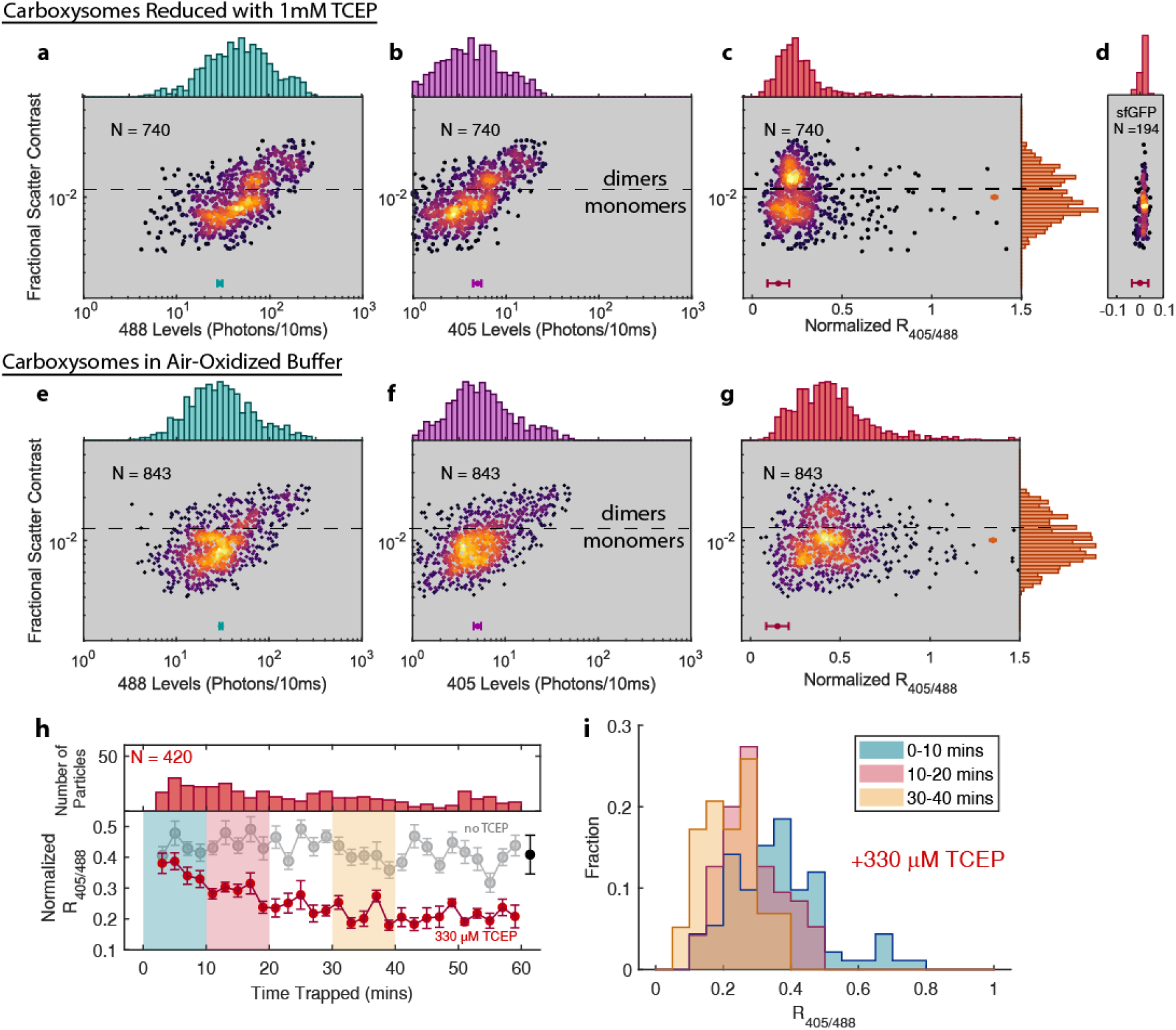
Multidimensional statistics of measurements on individual carboxysomes. (a)-(b) Scatter plots and marginal histogram between 488 and 405 nm fluorescence levels, respectively, with fractional scatter contrast, presented on logarithmic axes in both dimensions, due to the considerable range measured across carboxysomes. Each point corresponds to a single trapping event, colored by local density of points. The teal and magenta horizontal error bars denote the RMS standard error over all brightness levels. (c) Fractional scattering contrast *versus* ratiometric fluorescence *R*_405/488_ for carboxysomes in 1mM TCEP reducing buffer, with scattering contrast marginal histogram to the far right. Horizontal error bars denote the experimental RMS standard error in ratio uncertainty due to brightness fluctuations about each mean level (Supplementary Note S4 and Fig. S6). (d) The ratio-contrast scatter plot for sfGFP carboxysomes, demonstrating the narrow distribution measured for a reporter independent of redox. (e)-(g) Fluorescence, scatter contrast, and ratio scatter for carboxysomes in air-oxidized buffer, demonstrating ratios with a higher mean value and wider distribution than reduced carboxysomes. (h) Ratio kinetics measured after mixing with TCEP from individual carboxysomes. Red points correspond to ratios averaged for the number of carboxysomes shown in the top panel in 2 min windows after mixing with 330 μM TCEP, and gray points correspond to carboxysomes without TCEP reductant. Trapping starts approximately two minutes after mixing to load the sample into the cell, thus starting the experiment. Red and gray error bars indicate standard errors on the mean ratio measured in each time bin. Black error bar indicates the same RMS ratio error as in (g). (i) Ratio histograms from the reducing experiment, plotted in 10-minute intervals. At early times, the distribution is broad and centered at higher values, but gradually narrows and shifts to lower values over time.

Turning to redox ratios, Fig. 4c shows the 2D scatter plot correlating reduced roGFP2 fluorescence ratios *R*_405/488_ with the fractional scattering contrast of each carboxysome. The mean of the ratio distribution is 0.25 (*σ* = 0.16), a comparatively low value consistent with reducing conditions (Fig. S7). To test the ratio uncertainty due to measurement error, we also trapped *E*. *coli* carboxysomes labeled with superfolder GFP (sfGFP), whose ratiometric fluorescence is not redox-dependent (Fig. 4d, see also Fig. S8). The tight ratio distribution from sfGFP-labeled carboxysomes indicates that the larger ratio spread in roGFP2 carboxysomes arises from ratio variation between particles. To quantify the measured ratio uncertainty, we propagated the standard errors on the two fluorescence levels into their ratio (Note S4) and present the RMS standard error over all ratios (red error bars in Fig. 4c). In sfGFP carboxysomes, the ratio spread is comparable to the RMS uncertainty, indicating that measurement uncertainty dominates the spread. However, for reduced roGFP2 carboxysomes, the ratio distribution exceeds the bounds of the RMS error, indicating other contributions to the ratio spread, discussed further below. As well, the ratio distribution for roGFP2 narrows with larger scattering contrast, attributed to increased brightness from higher roGFP2 loading in some carboxysomes. While shot noise dictates that brighter fluorescence increases the absolute noise, the relative noise influencing ratio uncertainty is decreased.

When roGFP2-carboxysomes are left in air-oxidized buffer (Figs. 4e-4g), they display higher ratios on average (*μ* = 0.46) and a broader distribution (*σ* = 0.23). The mean ratio value is consistent with bulk redox ratio (Fig. S7), but the distribution shows an unexpectedly large spread. The RMS standard error on the ratios is comparable to the reduced case, but is distinctly smaller than the spread of the oxidized carboxysome ratio distribution, indicating additional heterogeneity. The ratio spread is insensitive to pH, added HCO_3_^−^, or added oxidant (1mM diamide, Fig S8). In particular, roGFP2-labeled carboxysomes are already in fully oxidizing environments when exposed to air. The wide spread of redox ratios likely arises from kinetics of individual roGFP2s which may occur on multiple timescales: sub-ms-timescale protonation/deprotonation kinetics,^40^ 10s of ms blinking into dark states,^41^ and the likely slower kinetics^42^ of the binding and unbinding of the engineered disulfide bridge on the roGFP2 β-barrel. The capabilities of the ISABEL trap combined with additional biological constructs will allow further investigation of the heterogeneity in oxidized samples in future work.

Figs. 4g and 4h demonstrate an hour-long reduction kinetics measurement of the ISABEL trap, where fluorescence ratios are measured on individual particles after mixing air-equilibrated samples into reducing buffer (330 μM TCEP). This measurement can be employed on dilute samples or where it is important to exclude ruptured fragments, large aggregates, and free roGFP2. Here, ratios are collected from individual carboxysomes as in Fig. 3 and averaged over 2-minute intervals, thus pooling roughly ~15 carboxysomes for each time point (numbers in upper panel of Fig. 4g). After mixing carboxysomes with reducing buffer, ratios decrease on a ~15-minute timescale (red trace in Fig. 4g) and settle at the reduced ratio mean of 0.22. This measurement recapitulates the ensemble reduction kinetics (Fig. S7), indicating that the bulk measurement is not dominated by external roGFP2. Along with the mean values in each time interval, the single-particle measurements allow us to measure the ratio distribution over time (Fig. 4h). In this case, the ratios initially show a broader spread, consistent with the oxidized population, which continuously shifts over time to the narrower distribution of reduced carboxysomes.

In summary, we demonstrated that individual carboxysomes can be trapped in solution with active feedback using interferometric detection of their optical scattering from a near-infrared laser. We introduced rapidly interleaving 405 and 488 nm excitation lasers to monitor the ratiometric fluorescence from individual roGFP2-labeled carboxysomes, decoupling fluorescence channels from the positional monitoring to measure intermittent signals with low excitation intensity. Carboxysomes recombinantly expressed in *E. coli* display wide distributions of scattering contrasts and fluorescence brightness, which can be directly monitored by trapping individual particles. Reduced and oxidized carboxysomes show low and high values of the average redox ratios *R*_405/488_, respectively. Controlling for chemical environment, single-carboxysome roGFP2 ratios display a wide range of values, indicative of the small numbers (*N* ≈ 3-15) of roGFP2 per carboxysome and other sources of heterogeneity, particularly evident in oxidized environments. We can observe minutes-timescale redox kinetics over the population of carboxysomes, which enables kinetic measurements for biological samples that are highly dilute or contain unwanted contributors to signal such as free roGFP2. Taken together, these experiments demonstrate the ability of the ISABEL trap to monitor nanoscale biological objects for extended times and to expand the range of local reporter experiments that can be done in these systems.

## Supplementary Material

See supplementary material for notes on sample preparation and characterization, ISABEL electronics and optics, and analysis and error notes. Supplementary figures show quantification of roGFP2 loading in carboxysomes, long trapping events, the impact of 405 nm illumination on ratiometric fluorescence, carboxysome sizing with cryo-EM, uncertainty statistics from trapping events, bulk reduction kinetic traces, 2D scatter plots for sfGFP carboxysomes, insensitivity of ratio distribution with buffer chemistry changes, and fluorescence spectra of roGFP2 and dye solution standards.

## Acknowledgments

This work was supported in part by the U. S. Department of Energy, Office of Science, Office of Basic Energy Sciences, Chemical Sciences, Geosciences, & Biosciences Division, Physical Biosciences Program, under Award Numbers DE-FG02-07ER15892 (W.E.M.) and DE-SC0016240 (D.F.S.). P.D.D. was supported in part by the Panofsky Fellowship at the SLAC National Accelerator Laboratory and by grant 2021-234593 from the Chan Zuckerberg Initiative DAF, an advised fund of Silicon Valley Community Foundation. D.D.P. was supported by a Stanford Graduate Fellowship.

## Data Availability

The data that support the findings of this study are available from the corresponding author upon request.

