## Supplemental Information for "Ratiometric Sensing of Redox Environments Inside Individual Carboxysomes Trapped in Solution"

### Supplementary Notes

#### Note S1: Microfluidic cell and sample preparation

We performed the trapping experiments in a quartz microfluidic cell that has been previously used successfully for ABEL and ISABEL experiments.[1, 2] The cells consist of two crossed channels that cross at a ~1.5-2 μm thin trapping region and four ports to load solutions and secure platinum electrodes for feedback in *x* and *y*. The top of the cell was chemically bonded with sodium silicate to a 0.15 mm thin quartz coverslip (Esco Optics). All measurements were made with the same cell, labeled C9.

Before each experiment, the internal surfaces of the cell were washed with 1M KOH, then passivated with a polyelectrolyte multilayer (PEM) consisting of four alternating layers of poly(ethylene imine) (PEI, Aldrich) and poly(acrylic acid) (PAA, Aldrich), serving as polycations and polyanions, respectively. The PEM multilayer sequence of PEI/PAA/PEI/PAA resulted in a uniform anionic surface which prevented nonspecific adhesion between anionic carboxysomes and the microfluidic cell via electrostatic repulsion. The layer deposition protocol was followed as previously described.[3]

After cell preparation, carboxysomes were diluted into HEPES buffer (10 mM HEPES, 15 mM NaCl, pH 7.5, 0.2 μm filtered) such that less than one carboxysome would be in the trapping area at any given time. Carboxysomes were stored at 4°C and were viable for up to 6 months after purification. For dilution, carboxysomes were drawn from the top of the stock suspension so as not to retrieve any large aggregates that had precipitated out of suspension. For the air-oxidized data in Figs. 4e-g in the main text, 0.5μL of purified N48-roGFP2-labeled *E. coli* carboxysomes with $A_{280}$ = 10.2 without $A_{340}$ baseline subtraction were diluted 700x into 350 μL of HEPES buffer, then gently mixed four times with ~ 150 μL of the sample volume to break up aggregates. For the trapping data in Figs. 4a-4c, carboxysomes were diluted 700x into HEPES buffer with 1mM TCEP and allowed to sit at room temperature for over an hour to allow the internal roGFP2 to fully reduce. sfGFP-labeled carboxysomes were diluted 200x in HEPES buffer and gently agitated prior to trapping. The concentration of the TCEP stock solution was confirmed with Ellman's reagent.[4]

For the kinetic reduction measurements, equal volumes of 350x-diluted roGFP2-labeled carboxysomes and a solution of 660 μM TCEP in HEPES were mixed then loaded into a cell such that trapping began within 2-3 minutes of mixing. Prior to mixing, the cell was quickly dried with $N_2$ and loaded onto the microscope stage centered on the trapping region. Feedback voltages were kept minimal to minimize anodic oxidation of TCEP at the electrodes.[5] Signatures of oxidized TCEP reaching the trapping area were detectable after about 40 minutes, as determined by monitoring the delayed fluorescence ratio rise of ~1 μM roGFP2 solution in 200 μM TCEP in the cell with feedback voltages applied on unlabeled polystyrene beads of size comparable to carboxysomes.

Between trapping runs, quartz cells were rinsed with Nanopure water, $N_2$ dried then cleaned overnight in piranha solution as described in Ref. 4.

#### Note S2: ISABEL Trap electronics and optics

<u>Near-IR Scatter Illumination</u>

The new trap design incorporates a near-IR diode laser centered at 802 nm to allow the high intensities typical of interferometric scattering experiments (>100 kW/cm$^2$)[6] while leaving the entire visible spectrum open for fluorescent channels without risk of photobleaching. The relay line for the scatter illumination beam is shown in Fig. S2. Illumination for scattering signal was provided by a multimode 802 nm laser diode (Axcel M9-808-150-D5P), driven by a Thorlabs LDC500 laser diode driver, housed in a

Thorlabs TCLDM9 mount, and temperature controlled by a Thorlabs TEC2000. To reduce the beam coherence, a bias tee in the diode mount was supplied with a sinusoidal AC wave from a function generator (Agilent 33220A) at 2.5MHz with $V_{pk\text{-}pk}$ = 2.00 V. With 50Ω impedance, this corresponds to a current modulation of 40 mA peak-peak on top of the 120 mA DC current from the laser diode driver.

The mode of the beam was circularized with a 4:1 anamorphic prism pair (Thorlabs) then focused into a single-mode fiber to produce ~ 15 mW of near-IR at the fiber output. The beam was polarized to horizontal polarization by a λ/2 plate, collimated and reduced to a spot size of $1/e^2$ diameter of 0.75 mm (via lenses $L_1$ and $L_2$ in Fig. S2) before guiding into the pair of AODs (AA Opto Electronic MT110-B50A1,5-IR). The x and y AODs deflect the beam in a predetermined, 32-point "Knight's tour" scan pattern[3] controlled by the field-programmable gate array (FPGA) on a National Instruments Reconfigurable Input/Output card (NI PCIe-7856R) via a direct digital synthesizer (AA Opto Electronic DDSPA2X-D4125b-34), described in more detail below. The first-order diffraction from each AOD was taken for generating the scan pattern. Lenses $L_3$ and $L_4$ map the pivot planes of AODx and AODy to each other, and were chosen to be the same focal length to ensure an equal aspect ratio of the scan pattern in the sample plane. Lenses $L_5$ and $L_6$ magnify the beam by 2 ($1/e^2$ diameter of 1.5 mm) and map the back focal plane (BFP) of the objective to the AODy pivot plane, to ensure purely angular changes in the BFP and thus lateral displacement in the sample plane. $L_5$ was placed 125 mm down the beam from AODy to allow 400 mm between $L_6$ and the BFP, ensuring enough room for the remaining optics in the illumination line.

The beam was then sent through a polarizing beam splitter (PBS), followed by a zero-order λ/4 plate, and reflected off a 775 nm shortpass dichroic mirror (Chroma), such that ~8 mW is focused by an NA 1.35, 100x oil immersion objective (Olympus UPlanApo Oil-Iris) to a 500 nm $1/e^2$ diameter spot in the sample plane. The beam is held at each position for 18.75 μs, such that the full 3×3 $\mu m^2$ area is scanned in 600 μs.

Scattering Detection

The reflected beam back-propagates through the quarter-wave plate, and is rotated to vertical polarization and reflected by the PBS to be collected on a photodiode (Newport 2031). The conjugate image plane at the photodiode is formed by focusing with a $f$ = 400 mm spherical lens after the PBS. An adjustable iris before the photodiode is used to allow only the image of the scan pattern to be incident on the photodiode. The photodiode is connected to a floating analog input channel on the FPGA. For each spot in the scan pattern, the FPGA waits 10 μs, then averages eight ADC-converted analog voltage measurements from the photodiode, each spaced 1 μs apart.

Feedback

The feedback scheme adopted here is the same as previously used.[2] Feedback voltages are applied by two pairs of platinum electrodes every 600 μs, determined by the detected position of the particle on the FPGA and amplified 8x by two op-amp circuits described in detail elsewhere.[7] For feedback, the position setpoint is specified to be near the center of the scan pattern, on a single scan point. Due to interactions between the buffer and the surface charges on the passivation layer, the applied electric fields steer the particle by electroosmosis. The feedback voltages can be tuned by gains ($g$ = 1.6 V/μm) and offsets (<160 mV) in homebuilt LabVIEW software. The FPGA calculates feedback voltage at each frame, and is linear with the displacement of the detected particle position from the trap setpoint in each dimension.

Fluorescence Excitation

The FPGA digitally modulates two collinear fluorescence excitation lasers every 1 ms for two-channel excitation of roGFP2. The 405 nm laser (Coherent Obis LX) is directly modulated by a digital output from the FPGA and spatially overlapped onto the path of the 488 laser beam (Coherent Sapphire), which is modulated via an AOD (Isomet 1205C-2, driver Isomet 222A-1) controlled by the FPGA. Both beams are circularly polarized by quarter-wave plates and attenuated such that ~30-40 μW in each beam reach the sample plane. The collinear beams are focused with an $f$ = 400 mm Köhler lens and reflected off a multi-band dichroic (Semrock Di03-R405/488/561/635-t1-25x36) for wide-field illumination in the sample plane ($1/e^2$ radius ≈ 4 μm). Low intensity (< 50 $W/cm^2$) is necessary at 405 nm to reduce the probability of excited state proton transfer[1] and photoconversion of roGFP2 chromophores over extended trapping times (Fig. S3). For these experiments, the peak intensities of the 488 and 405 nm excitation beams are 80 and 40 $W/cm^2$, respectively.

Fluorescence Detection

Emitted fluorescence is spectrally separated from the scatter illumination and fluorescence excitation beams by sequential transmission through the 775 nm shortpass and multi-bandpass dichroic mirrors specified above. The fluorescence is spatially filtered by a 75 μm-diameter pinhole centered on the trapping center position (corresponding to 1.5 μm diameter at the sample) and spectrally filtered to collect the 500-570 nm emission band of roGFP2 (Fig. S9) before detection on an avalanche photodiode (PicoQuant τ-SPAD) connected to the FPGA. To distinguish photons emitted from 405 and 488 nm excitation, each time-tagged photon is additionally tagged with the identity of the excitation laser for separation of the two channels in post-processing.

Control

All of the components above are controlled and synchronized by an FPGA on a reconfigurable input-output board (NI PCIe 7856) with an 80 MHz clock and custom software written in LabVIEW, as previously implemented.[2] This control allows for the calculation of absolute fractional scattering contrast for each 600 μs frame. The FPGA then calculates and applies voltages in x and y that is proportional to the displacement of the maximum absolute fractional scattering contrast to the feedback setpoint.

#### Note S3: Analysis

Calculation of Interferometric Scattering Contrast

Absolute fractional scattering contrast describe in Eq. 2 of the main text is calculated in real time on the FPGA for particle localization, described in detail previously.[2] Before the measurement, we measure the ADC counts on the detector with the near-IR beam blocked as an offset to subtract from the measured ADC counts at each point. During the experiment, a 10-ms average of the ADC counts at each scan point is taken when feedback is turned off to provide the reference background intensity $|E_r|^2$ in Eq. 2. Subsequent measurements are divided by this background at that scan point to help identify the pixel of maximum fractional scattering contrast over each set of 32 beam-dwell times in a "frame". To account for small power fluctuations in laser power between each 600 ms frame, the average of the ADC values of the outermost 16 scan points are used to normalize each background measurement.[2] For post-processing and analysis, the ADC counts (averaged from 8 consecutive measurements as described above) from each point and their corresponding background measurements are saved. The absolute fractional scattering contrast for offline analysis plotted in Fig. 3 of the main text and used in level-finding is determined from the maximum value within a 3x3 pixel area around the trap setpoint.

Determination of fluorescence and scattering contrast levels

Mean brightness and scattering contrast levels from each trapping event are determined by a changepoint-finding algorithm[8] on the 488 nm level trace due to its highest signal-to-background ratio. Level values were calculated as mean values between two changepoint times. The determined changepoint times were used on the 405 nm and scattering contrast traces to determine their average values in the same time intervals. These levels can consist of trapping events or times with only background.

The determined levels are filtered and processed to ensure each point in the 2D scatter plots of Fig. 4 of the main text correspond to trapped carboxysomes. Levels that meet all of the following criteria are kept for further analysis, while levels that do not meet the criteria are rejected. Admissible levels:

- Only occur when feedback is ON, to remove stuck particles that persist when feedback is OFF.
- Only occur when feedback voltage polarity is correct, to make sure all particles have the same sign on charge.
- Span at least 200 ms, so that particles diffusing in the ROI without feedback are ignored.
- Have brightness in the 488 channel exceeding $2\sigma$ from the mean of background levels, determined as described below.
- Have brightness in the 405 channel at least 1 count/10 ms above background.
- Have absolute fractional scattering contrast between 0.0025 and 0.025, to ignore small particles and large aggregates that do not correspond to single or double carboxysomes.

Fluorescence background values for each level in the 488 and 405 nm channels are determined as the mean of the fluorescence levels during the trap OFF times immediately before and after the trapping event. These background levels are taken as their median fluorescence to mitigate the influence of fluorescence bursts from rapidly diffusing objects.

We also merge consecutive levels with absolute fractional scattering contrasts that are within 10% of each other, since fluorescence dynamics such as blinking in the 488 nm channel may give the false appearance of separate levels within a single trapping event.

In summary, particles saved for analysis reflect objects that are trapped for at least 200 ms and show scattering contrasts in an appropriate range and also show fluorescence brightness in both 405 and 488 channels. These filters are reflected in the traces shown in Fig. 4 of the main text and are used to determine the fluorescence ratios from each particle.

Determination of fluorescence ratios

Ratios from each trapping event are determined via the following expression

$$R_{405/488} = \frac{I_{405} - b_{405}}{I_{488} - b_{488}}, \quad \text{(S1)}$$

where $I_{405}$ and $I_{488}$ refer to the mean fluorescence brightness over the trapping event in the 405 and 488 nm channels respectively, and $b_{405}$ and $b_{488}$ refer to the background levels determined for each level. To account for day-to-day changes in pointing of the fluorescence excitation beams, we normalize each ratio to the ratiometric fluorescence from Rhodamine 110 in water ($R_{min}$) and Coumarin 6 in absolute ethanol ($R_{max}$), measured after each trapping experiment, whose excitation spectra are shown in Fig. S10. For the 405 nm and 488 nm intensities used in this experiment, $R_{min} \approx 0.02$ and $R_{max} \approx 0.45$. Ratio normalizations are calculated according the following expression:

$$R_{norm} = \frac{R_{405/488} - R_{min}}{R_{max} - R_{min}}, \quad \text{(S2)}$$

Such that the ratio values from the Rhodamine 110 and Coumarin 6 standards are 0 and 1, respectively. We initially attempted ratio normalizations with roGFP2 pushed to oxidized and reduced extremes with 1 mM $H_2O_2$ and 1 mM TCEP, respectively, but found that the obtained ratio values were not as reliable. This may have arisen from unspecific adsorption of roGFP2 to the microfluidic cell surfaces in a way that impacted the protonation equilibrium of the roGFP2 chromophore.

#### Note S4: Uncertainty Estimation

Standard errors on the two fluorescence levels and scattering contrast were determined for each trapping event via a bootstrapping function in MATLAB. For each level, 100 bootstrapped means were determined from the values within a level by sampling with replacement, and the standard error on the mean (SEM) was taken as the standard deviation on the distribution of bootstrapped means. We compare the error statistics for 488 levels, 405 levels, and ratios from oxidized roGFP2-carboxysomes, reduced roGFP2-carboxysomes, and sfGFP carboxysomes in Tables S1-S3 below. As shown in Fig. S8a-f, the uncertainties in air-oxidized carboxysomes were compared to the expected SEM with Poisson statistics (assuming a level with a mean bright *I* plus background *b*), as an "excess noise ratio" (measured SEM/shot noise SEM). For example, the RMS excess noise ratio on the 488 levels in reduced carboxysomes was 1.52x the expected value for shot noise. By contrast, the RMS excess noise ratio in the 405 channel was only 1.04, indicating closer agreement with the shot noise limit.

The SEMs and mean values were used to propagate the uncertainty to ratio measurements:

$$\left(\frac{SEM_R}{R}\right)^2 = \left(\frac{SEM_{405}}{I_{405}}\right)^2 + \left(\frac{SEM_{488}}{I_{488}}\right)^2. \quad \text{(S3)}$$

In Figs. S7g-S7i, the fractional uncertainty on the 405 channel levels are higher due to the lower brightness and larger contribution from background. The fractional uncertainty of the ratio measurements is therefore dominated by the uncertainty in the level of the 405 nm channel.

For the normalized ratios presented in the main text, the fractional uncertainties were scaled accordingly. From Eq. S2, the standard error $SEM_{norm}$ on the normalized ratio $R_{norm}$, can be calculated as follows:

$$\frac{SEM_{norm}}{R_{norm}} = \frac{SEM_R}{R_{max} - R_{min}} \cdot \left(\frac{R - R_{min}}{R_{max} - R_{min}}\right)^{-1} = \frac{SEM_R}{R - R_{min}}. \quad \text{(S4)}$$

Table S1: Error Statistics for oxidized roGFP2-carboxysomes

| Channel | Mean | RMS SEM | RMS Fractional error | RMS Excess Noise Ratio |
|---|---|---|---|---|
| 488 nm | 39.9 cts/10ms | 1.0 cts/10ms | 0.04 | 1.37 |
| 405 nm | 7.3 cts/10ms | 0.5 cts/10ms | 0.12 | 1.05 |
| Norm. $R_{405/488}$ | 0.45 | 0.06 | 0.12 | 1.08 |

Table S2: Error Statistics for reduced roGFP2-carboxysomes

| Channel | Mean | RMS SEM | RMS Fractional error | RMS Excess Noise Ratio |
|---|---|---|---|---|
| 488 nm | 57.6 cts/10ms | 1.6 cts/10ms | 0.04 | 1.50 |
| 405 nm | 5.3 cts/10ms | 0.5 cts/10ms | 0.14 | 1.03 |
| Norm. $R_{405/488}$ | 0.29 | 0.06 | 0.18 | 1.05 |

Table S3: Error Statistics for sfGFP-carboxysomes

| Channel | Mean | RMS SEM | RMS Fractional error | RMS Excess Noise Ratio |
|---|---|---|---|---|
| 488 nm | 583.43 cts/10ms | 7.0 cts/10ms | 0.01 | 2.11 |
| 405 nm | 42.3 cts/10ms | 1.0 cts/10ms | 0.02 | 1.11 |
| Norm. $R_{405/488}$ | 0.01 | $7\times10^{-4}$ | 0.07 | 1.19 |

#### Note S5: Growth and purification of labeled *E. coli* carboxysomes

*E. coli BW25113* harboring a plasmid containing the *H. neapolitanus* HnCB10 carboxysome operon from Bonacci et al.[9] and a plasmid containing roGFP2 with an N-terminal carbonic anhydrase "N48" tag (first 53 amino acids of CsosCA) were inoculated into 1 L of LB at half concentrations of antibiotic on each plasmid. Cells were grown to log phase and induced with 500 μM IPTG and 100 nM aTc, then grown overnight at 18°C before pelleting and freezing at -20°C. Frozen pellets were lysed in BPER reagent with the addition of 10 mM $MgCl_2$, 20 mM $NaHCO_3$, 1 mM EDTA, 0.1 mg/mL lysozyme, 1 mM PMSF, and 1 μL benzonase/25 mL lysate. Lysis occurred for 1 hour on a shaking rotor at room temperature. To clarify, cells were spun at 12,000 xG for 15 minutes, and the supernatants spun again at 40,000 xG for 30 minutes to pellet carboxysomes. The pellets were resuspended in 1.5 ml of TEMB buffer (10 mM Tris-HCl, 10 mM $MgCl_2$, 20 mM $NaHCO_3$, 1 mM EDTA, adjusted to pH 8) and gently shaken on ice from 30 minutes to overnight to loosen the pellet before resuspending. Resuspended pellets were spun at 900 xG for 3 minutes to pellet insoluble junk before loading onto a 25 ml 10-50% sucrose gradient. The sucrose gradient was spun at 105,000 xG for 35 minutes and 1 ml fractions collected and analyzed on SDS-PAGE. Carboxysome fractions were pooled and spun at 100,000 xG for 90 minutes, the supernatant dumped, and pellets resuspended in 250 ul of TEMB and stored at 4°C. The final concentration of roGFP2 labeled carboxysomes as determined by absorbance at 280nm was 10.2. The $A_{280}$ reported here is without 340 nm baseline subtraction because carboxysomes contribute to baseline scattering. Superfolder-GFP *E. coli* carboxysomes with FLAG tag on the N-terminus of CsoS1A were prepared and purified in the same was as roGFP2-labeled *E. coli* carboxysomes. Carboxysomes were stored at 4°C and were viable for up to 6 months after purification.

#### Note S6: Quantifying roGFP2 brightness and roGFP2 loading per carboxysome

Copy numbers of active roGFP2-labeled carboxysomes were determined by analyzing bleachdown traces for single-step roGFP2 photobleaching. roGFP2-labeled carboxysomes were diluted 1200x in HEPES buffer and reduced for 1 hour with 1 mM TCEP, so as to push the equilibrium to the 488 nmabsorbing anionic chromophore. The reduced carboxysomes were then introduced into a hybridization cell (Grace Bio-Labs Secure Seal) stuck to a quartz coverslip (Esco Optics) and coated with PEI/PAA/PEI as described in Note S1. The polycation PEI served as the final layer to encourage electrostatic adsorption to the coverslip surface. Carboxysomes were then illuminated with 488 nm (240 $W/cm^2$ peak intensity, $1/e^2$ radius ~ 4μm) and imaged at 20 fps on an sCMOS camera (Andor Zyla 4.2) for 2 minutes to allow for complete photobleaching. The camera was calibrated[10] to convert ADU counts to photons. Time traces were constructed from the 5x5 pixel ROI (875x875 $nm^2$) to collect all detected photons around each carboxysome point-spread function. Brightness step sizes due to photobleaching or blinking were determined via a changepoint-finding algorithm[8] on the time traces, then manually examined to remove spurious step changes. The illumination profile was also measured to normalize measured brightness steps and exclude carboxysomes where excitation intensity was less than 1/e of the peak. Normalized brightness steps were collected to give the histogram in Fig. S1a. Because multiple steps may occur within the 50 ms frame, there is an expected tail to higher brightnesses. Total carboxysome loading (Fig. S1b) was estimated by taking the ratio of the initial brightness level determined by the changepoint algorithm over the median single brightness step size, normalized by excitation beam intensity. Most carboxysomes appear to contain 3-15 active roGFP2 molecules, but we also observe a tail to higher loading.

#### Note S7: Imaging and Sizing Carboxysomes with Cryo-Electron Microscopy

3 μL of carboxysome suspension (diluted 10x in HEPES buffer from stock) was deposited onto a glow-discharged holey carbon electron microscopy grid (Quantafoil R 2/2 G200F1), blotted on both sides for 2.5 seconds, and plunge frozen (Gatan CP3). Electron micrographs were acquired on a 200-keV electron microscope (Thermo Fisher Glacios) equipped with a direct detector (Gatan K2). Images were acquired with pixel spacing of 2.43 Å and 12.7 Å, with defocus targets of -3 μm and -40 μm respectively.

Effective diameters were determined by measuring the area of the polygon traced around the carboxysome shells in Fiji,[11] then evaluated as $d_{eff} = (4A/\pi)^{1/2}$ assuming circular shape. As shown in Fig. S5, the diameters appear normally distributed with $\mu$ = 141 nm and $\sigma$ = 31 nm. This is in contrast to the bimodal scattering contrast histograms shown in Fig. 4 of the main text, strongly suggesting that the larger-contrast feature is due to carboxysome dimers. The distribution of effective radii (coefficient of variation $\sigma/\mu$ = 0.22) also results in a spread of volumes, which generates substantial overlap between the single- and double-carboxysome features in the scattering contrast histogram.

#### Supplementary Figures

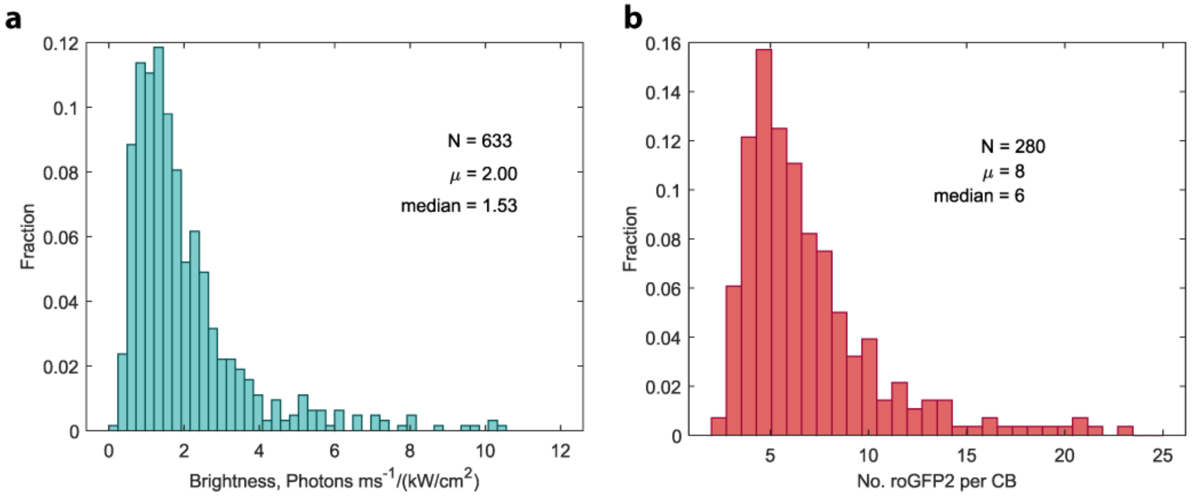

**Figure S1.** Quantifying roGFP2 single-molecule brightness and roGFP2 loading in individual carboxysomes. (a) Distribution of roGFP2 brightnesses measured via single-step photobleaching events in reduced carboxysomes, described in detail in Note S6 above. This histogram primarily reports on the bleaching of single roGFP2 copies, but the possible simultaneous bleaching of two or three copies are also incorporated into this histogram. (b) Estimation of number of roGFP2 copies per carboxysome, determined by dividing the initial brightness of each carboxysome by the median single-roGFP2 value measured in (a).

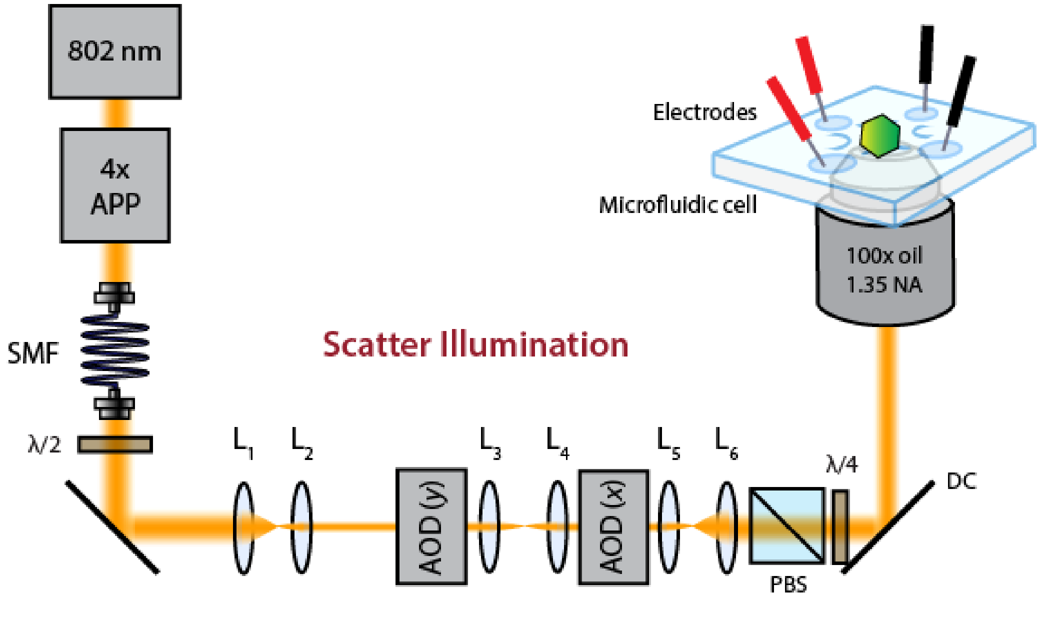

**Figure S2.** Scatter illumination relay line, described in detail in Note S2. 4x APP: 4x anamorphic prism pair. SMF: single-mode fiber. λ/2: half-wave plate. $L_1$-$L_6$: spherical lenses with focal lengths $f_1$ = 100 mm, $f_2$ = 50 mm, $f_3$ = 60 mm, $f_4$ = 60 mm, $f_5$ = 150 mm, $f_6$ = 300 mm. AOD: acousto-optic deflector. PBS: polarizing beam splitter. λ/4: quarter-wave plate. DC: dichroic mirror.

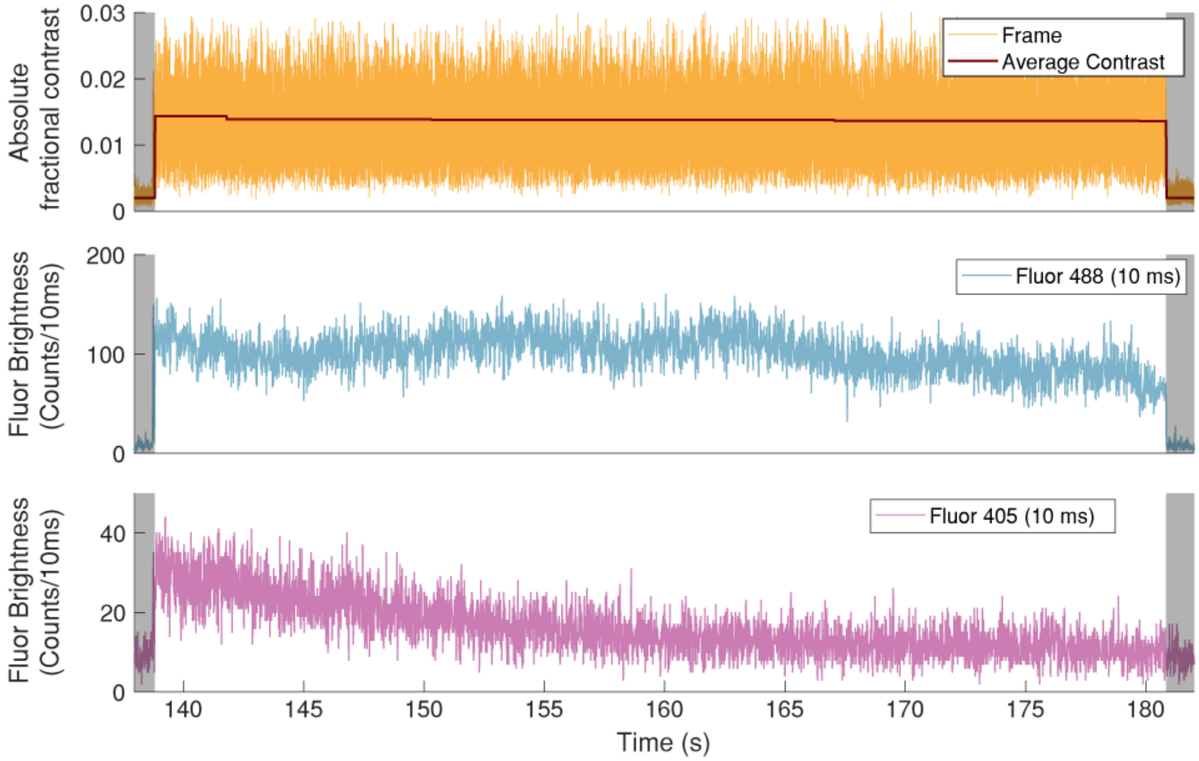

**Figure S3.** Carboxysomes can be trapped for more than 40 seconds via localization with interferometric scattering. Top trace: absolute fractional scattering contrast remains level while trapping feedback is ON. Middle trace: 488 nm-excited fluorescence trace shows a consistent brightness over the whole trapping event. Bottom trace: 405 nm excitation fluorescence shows a gradual reduction in brightness due to excited state proton transfer (ESPT)[12] from the chromophore.

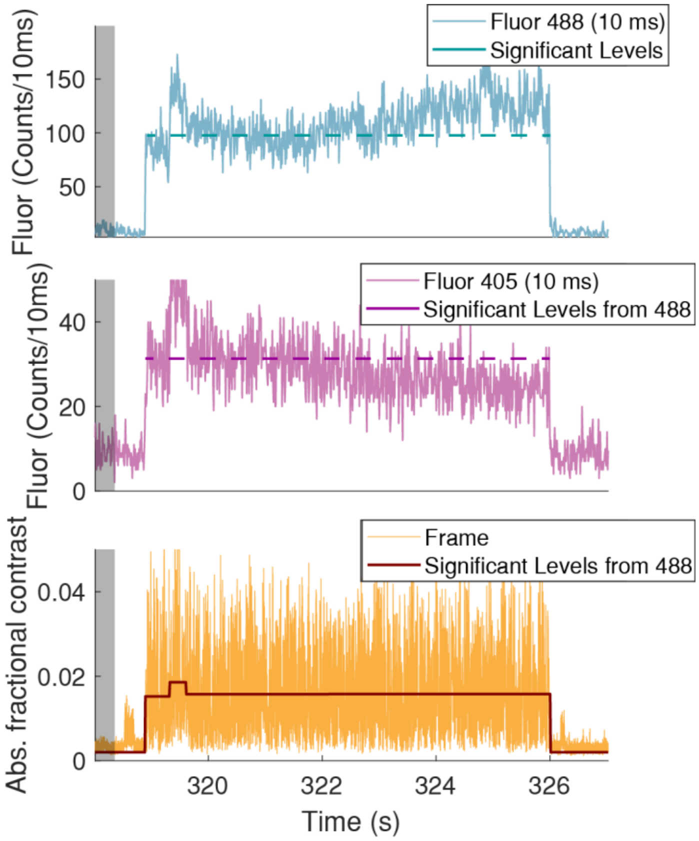

**Figure S4.** An example of 405 nm-induced excited state proton transfer (ESPT) introducing slow drifts into fluorescence levels. Top and middle traces: 488 fluorescence brightness increases and 405 fluorescence decreases as the roGFP2 chromophore converts to its anionic form. Dashed lines indicates mean fluorescence brightness levels from around $t \approx 320$ s. Bottom trace: absolute fractional scatter trace with average value. From $t = 320$ s onwards, the scatter contrasts remains level, indicating the same particle is trapped until $t = 326$ s.

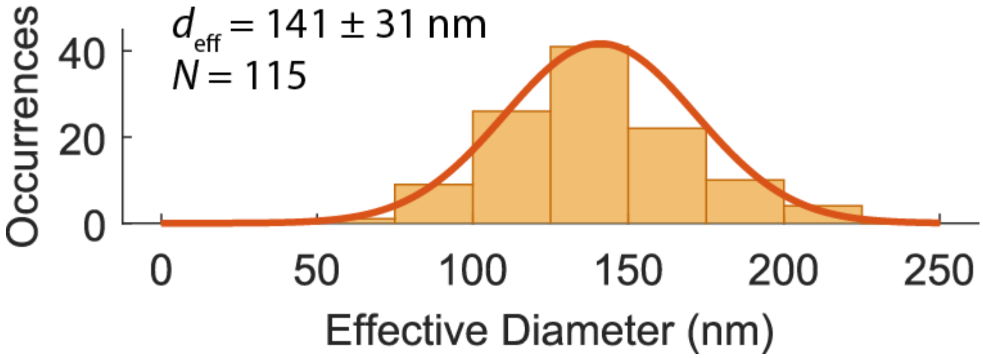

**Figure S5.** Distribution of effective diameters of roGFP2-labeled *E. coli* carboxysomes determined with cryo-EM imaging (see Note S7). Error on $d_{eff}$ denotes standard deviation. Dark orange line shows a Gaussian fit to the histogram.

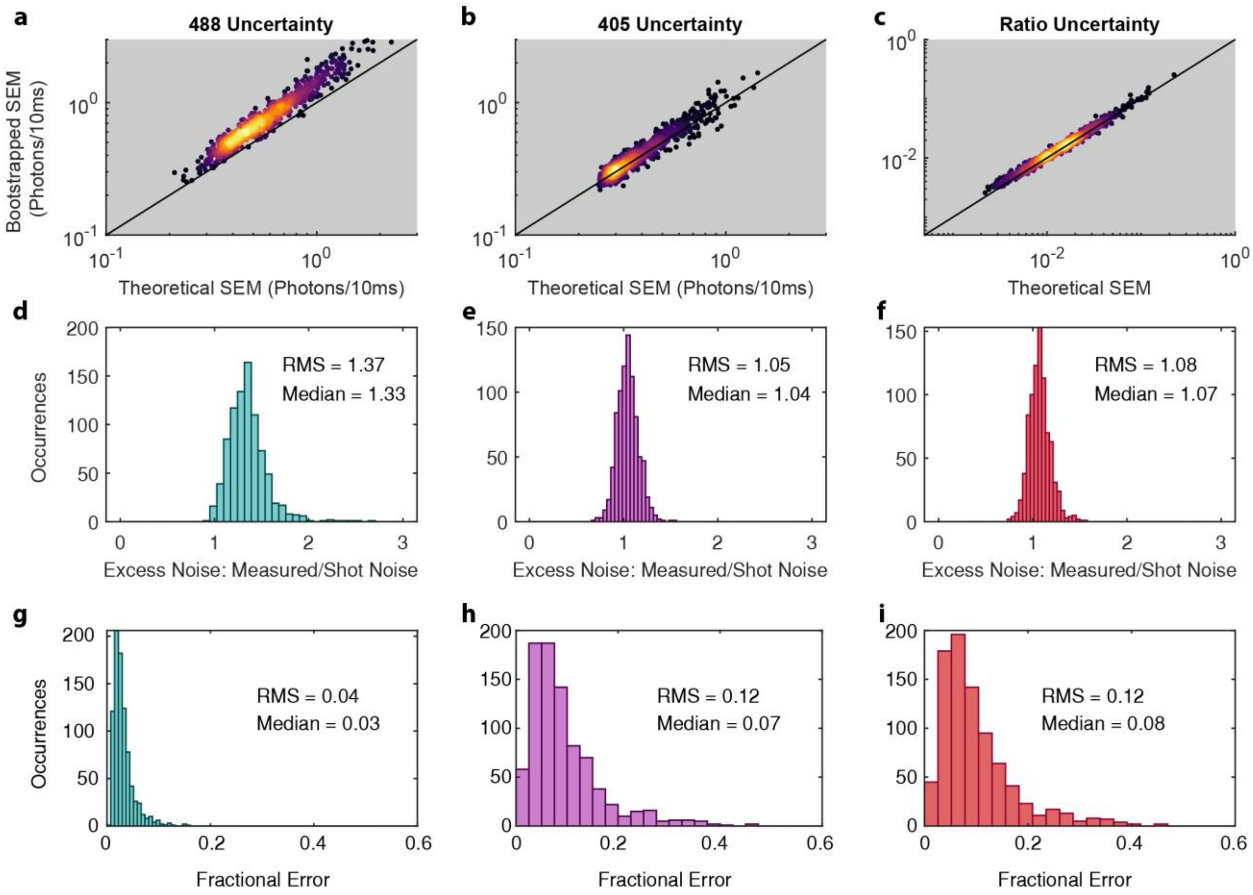

**Figure S6**. Statistics of standard errors on the mean (SEM) of each brightness level for carboxysomes in air-oxidized buffer. Derivations of the quantities plotted are presented in Supplemental Note S4. (a)-(c) Comparison of the theoretical and measured SEMs on 488 brightness, 405 brightness, and ratio, respectively. Theoretical SEM is based on expected uncertainty arising from shot noise, while measured SEMs are derived from bootstrapping from the brightness values within each level. The black diagonal line represents when the measured noise is equal to shot noise. Ratio SEMs in both dimensions are determined by error propagation on the 488 and 405 nm SEMs. (d)-(f) Histograms of the excess noise ratio of measured and shot noise SEMs on each level for 488, 405, and ratio uncertainties, respectively. (g)-(i) Fractional uncertainties SEM/$\mu$ for 488, 405, and ratio levels. The bottom row demonstrates that the biggest source of uncertainty in the ratio measurement arises from noise on the 405 levels. This arises from the lower brightnesses and higher backgrounds on these levels compared to 488 nm.

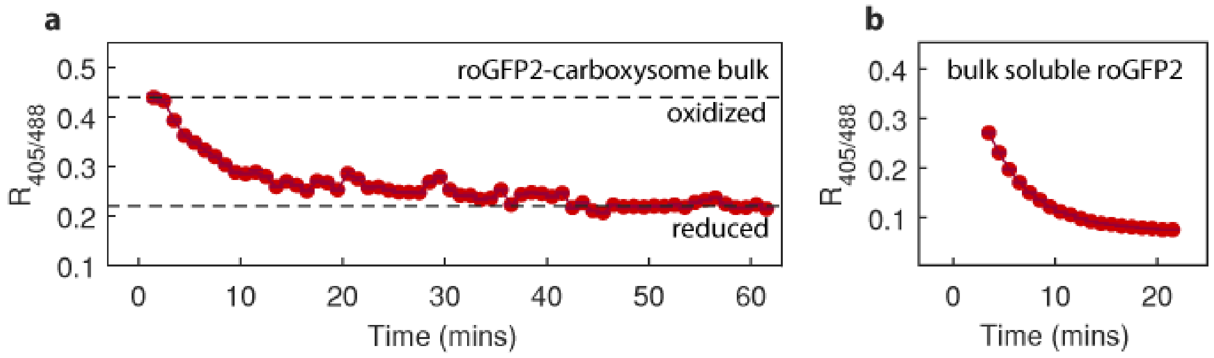

**Figure S7**. Ratiometric fluorescence kinetics from bulk samples after adding 330 µM TCEP: (a) roGFP2-labeled carboxysomes and (b) soluble 0.5 µM N48-roGFP2. The dashed lines in (a) show the typical ratio values in oxidizing and reduced conditions, consistent with the mean ratios measured from single carboxysomes. Ratio measurements start after approximately 2 minutes due to time needed for sample loading after mixing. Soluble roGFP2 reduces ratio more quickly in free solution and reduce to lower ratio values than roGFP2-labeled carboxysomes.

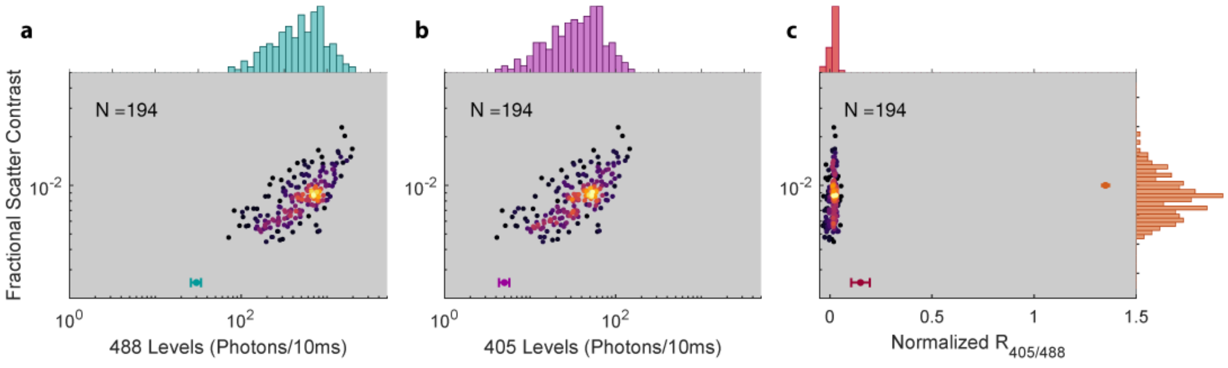

**Figure S8**. Scatter plots relating brightnesses, scatter contrast, and fluorescence ratio in trapped sfGFP carboxysomes.

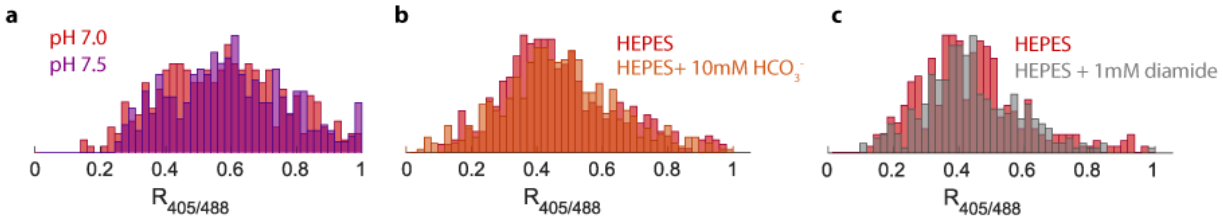

**Figure S9**. Changes in buffer chemistry do not significantly impact the wide ratio spread in air-oxidized carboxysomes. (a) Comparison between trapped carboxysomes in pH 7.0 and pH 7.5 citrate-phosphate buffer. (b) Comparison between trapped carboxysomes in HEPES buffer with or without 10 mM sodium bicarbonate. (c) Comparison between trapped carboxysomes in HEPES buffer with or without 1 mM diamide. This test confirms that air-oxidized carboxysomes are indeed already fully oxidized. In (b) and (c), HEPES buffer refers to 10 mM HEPES + 15 mM NaCl, pH 7.5 as used in trapping experiments.

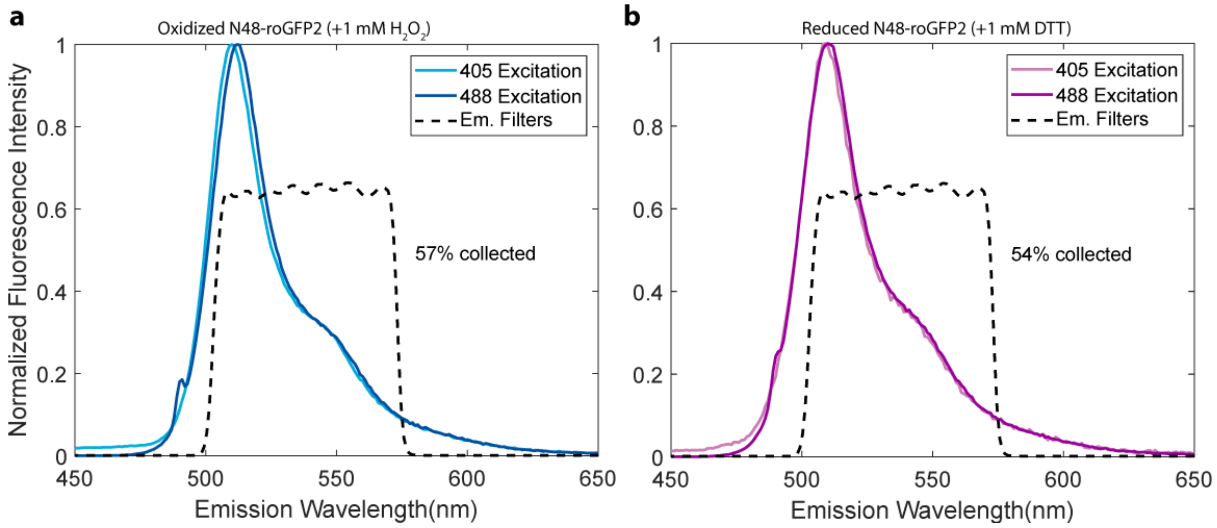

**Figure S10**. Emission spectra from 1μM N48-roGFP2 in Tris buffer when pushed to (a) full oxidation or (b) full reduction. The peak position can shift by a few nm, but the whole emission spectrum is collected by our 500-570 nm emission filter set regardless of redox state. Filter set: Chroma HQ500 longpass, Semrock EdgeBasic 488 longpass, 570 shortpass, 785 shortpass, 808 notch.

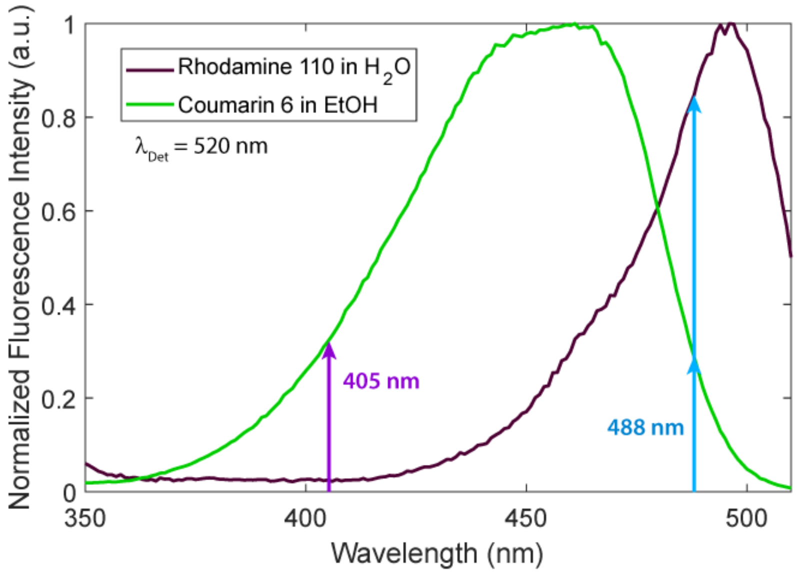

**Figure S11**. Fluorescence excitation spectra of Rhodamine 110 and Coumarin 6 used as standards for normalizing measured roGFP2 ratios to low and high values, respectively. See Note S3 for ratio normalization.
